# Contraception ends the genetic maintenance of human same-sex sexual behavior

**DOI:** 10.1101/2023.03.07.531528

**Authors:** Siliang Song, Jianzhi Zhang

## Abstract

Because human same-sex sexual behavior (SSB) is heritable and leads to fewer offspring, it is puzzling why SSB-associated alleles have not been selectively purged. Current evidence supports the antagonistic pleiotropy hypothesis that SSB-associated alleles benefit individuals exclusively performing opposite-sex sexual behavior by increasing their number of sexual partners and consequently their number of offspring. However, here we show that having more sexual partners no longer predicts more offspring since the availability of oral contraceptives in the 1960s and that SSB is now negatively genetically correlated with the number of offspring, indicating a loss of SSB’s genetic maintenance in modern societies.

## Introduction

About 2-10% of individuals across human societies perform same-sex sexual behavior (SSB) (1), a trait with a broad-sense heritability of ~30% (1, 2). We refer to those who perform SSB as SSB individuals, regardless of whether they perform SSB exclusively. We refer to those who perform only opposite-sex sexual behavior (OSB) as OSB individuals. Because SSB individuals have substantially lower numbers of offspring than OSB individuals (1), one wonders why SSB-associated alleles have not been selectively purged in evolution (3). A recent study discovered a strong positive genetic correlation between SSB and the number of sexual partners among OSB individuals (4), meaning that OSB individuals carrying SSB-associated alleles have more sexual partners than those not carrying such alleles. It is generally believed that, under polygyny, which was common in human history, a male with more sexual partners typically has more children (5, 6). There is also evidence from traditional human societies that the number of sexual partners of a female positively predicts her number of children (7). Therefore, the above discovery suggests that SSB-associated alleles offer a reproductive advantage to OSB individuals at least in premodern societies, which could in principle explain the evolutionary maintenance of these alleles under the general hypothesis of antagonistic pleiotropy (4).

However, rapid societal changes in the last 50-100 years have greatly shifted attitudes toward and practice of reproduction (8, 9). In particular, the modernization and popularization of contraception since the 1960s may have largely decoupled the number of sexual partners of OSB individuals from their number of offspring (10), which could abolish the aforementioned mechanism for the genetic maintenance of SSB. To test this hypothesis, we used the UK Biobank (UKB), a database of genetic and phenotypic information of about 0.5 million British participants who were born between 1937 and 1970 and recruited between 2006 and 2010 (11).

## Results and Discussion

We first investigated whether the number of sexual partners and number of children are correlated phenotypically among OSB individuals. To prevent population stratification, we focused on UKB participants with European ancestry—the largest ethnic group in the UKB (*N* = 452,557), from which 374,585 participants answered questions on sexual partners and children; these answers allowed the identification of 358,861 OSB individuals. Phenotypic regression of the number of children on the number of sexual partners was performed with covariates of genetic sex, age, age squared, and the first ten genetic principal components. Socioeconomic confounding factors were also controlled by including household income, deprivation index, number of years of education, year of first sexual intercourse, and their squared terms in the covariates. When males and females were analyzed together, the number of sexual partners was found to negatively predict the number of children among OSB individuals (coefficient *β* = - 2.3×10^-3^ *P* = 1.6×10^-90^; **Table 1**). A much stronger effect was observed among females (*β* = - 1.5×10^-2^, *P* < 2.2×10^-308^) than among males (*β* = −7.2×10^-4^, *P* = 2.7×10^-9^).

**Table 1.**
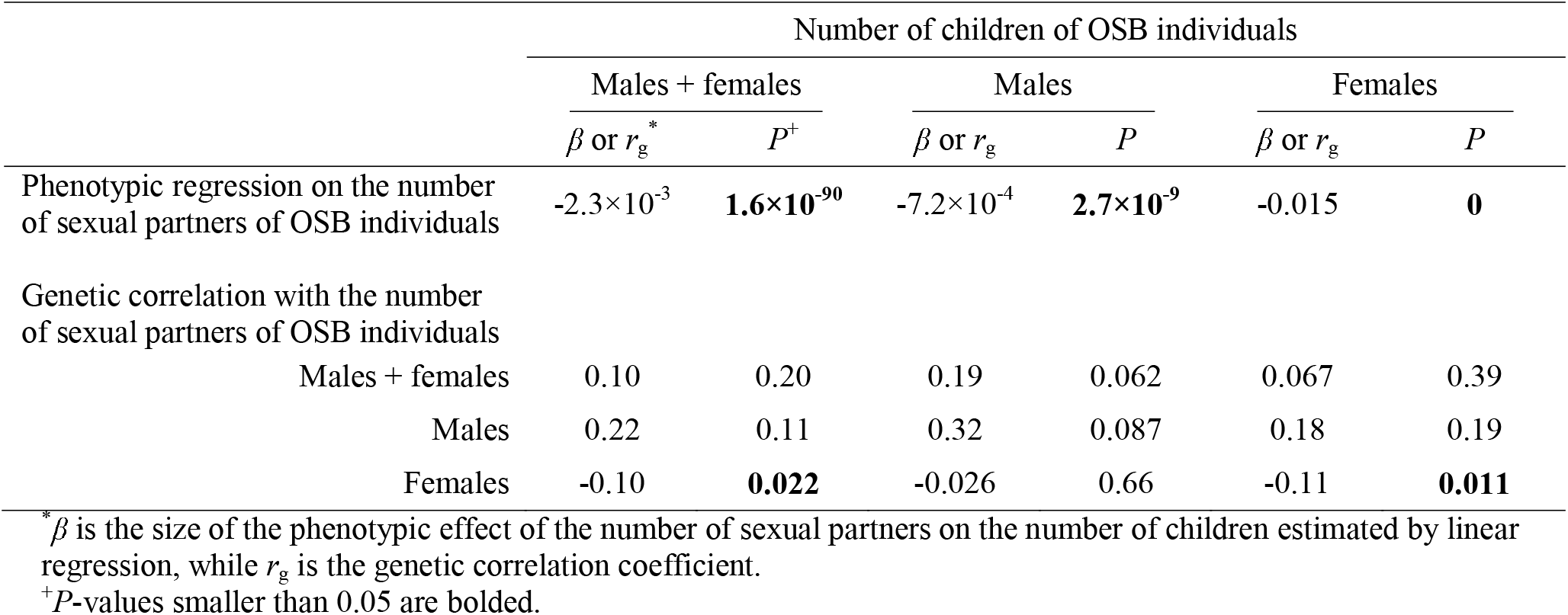
Phenotypic and genetic correlations between the number of sexual partners and the number of children in OSB individuals

We then estimated among OSB individuals the genetic correlation between the number of sexual partners and the number of children, which measures the proportion of variance that the two traits share due to genetic causes. Genome-wide association analysis (GWAS) was respectively performed on these two traits in the OSB European ancestry population mentioned to obtain summary statistics (see Materials and Methods), based on which the genetic correlation was computed using cross-trait LD score regression (12). The genetic correlation is not statistically significant in male and female OSB individuals analyzed together (*r*_g_ = 0.10, *P* = 0.20), nor in male OSB individuals (*r*_g_ = 0.32, *P* = 0.09), but is significantly negative in female OSB individuals (*r*_g_ = −0.11, *P* = 0.01) (**Table 1**).

Generally consistent patterns were respectively observed for the above phenotypic (**Dataset S1**) and genetic (**Dataset S2**) correlations when the samples from 19 UKB recruitment centers (>10,000 participates per center) broadly distributed across the UK were individually analyzed, suggesting that our results are general. These findings suggest that, contrary to the expectation from traditional societies (5–7), the number of sexual partners of OSB individuals does not phenotypically or genetically positively predict their number of offspring in the contemporary British population (**Table 1**). Consequently, the previously reported strong positive genetic correlation between SSB and the number of sexual partners among OSB individuals (4), which we replicated in the UKB (**Dataset S4**), does not translate to a positive genetic correlation between SSB and the number of children among OSB individuals (*r*_g_ = −0.027, *P* = 0.61) (**Dataset S5**).

A substantial increase in contraception took place in the UK after oral contraceptives became available to married and unmarried women in 1961 and 1967, respectively (13). To more specifically evaluate the potential role of contraception in the above finding, we divided the UKB data into cohorts in subsequent analysis. Because >99% of UKB participants had their first sexual intercourse between 1950 and 1989 (**Dataset S3**), we divided all OSB participants into eight 5-year cohorts according to their year of first sexual intercourse. We then examined the phenotypic effect of the number of sexual partners on the number of children in each cohort by regression (see Materials and Methods). Interestingly, the coefficient *β* was positive in the first two cohorts (1950-1959) but became negative in all subsequent cohorts (1960-1989) (**Fig. 1**). The switch of the sign of *β* coincided with the availability of oral contraceptives and popularization of contraception in the UK. Because the 1950-1954 cohort were most likely still sexually active after 1960, their *β* value may have also been impacted by increased contraception. Hence, although the UKB does not allow analyze sufficient individuals who became sexually active before 1950, their *β* was likely even more positive than that for the 1950-1954 cohort, supporting the previously proposed mechanism of the genetic maintenance of SSB in premodern societies (4).

**Fig. 1.**
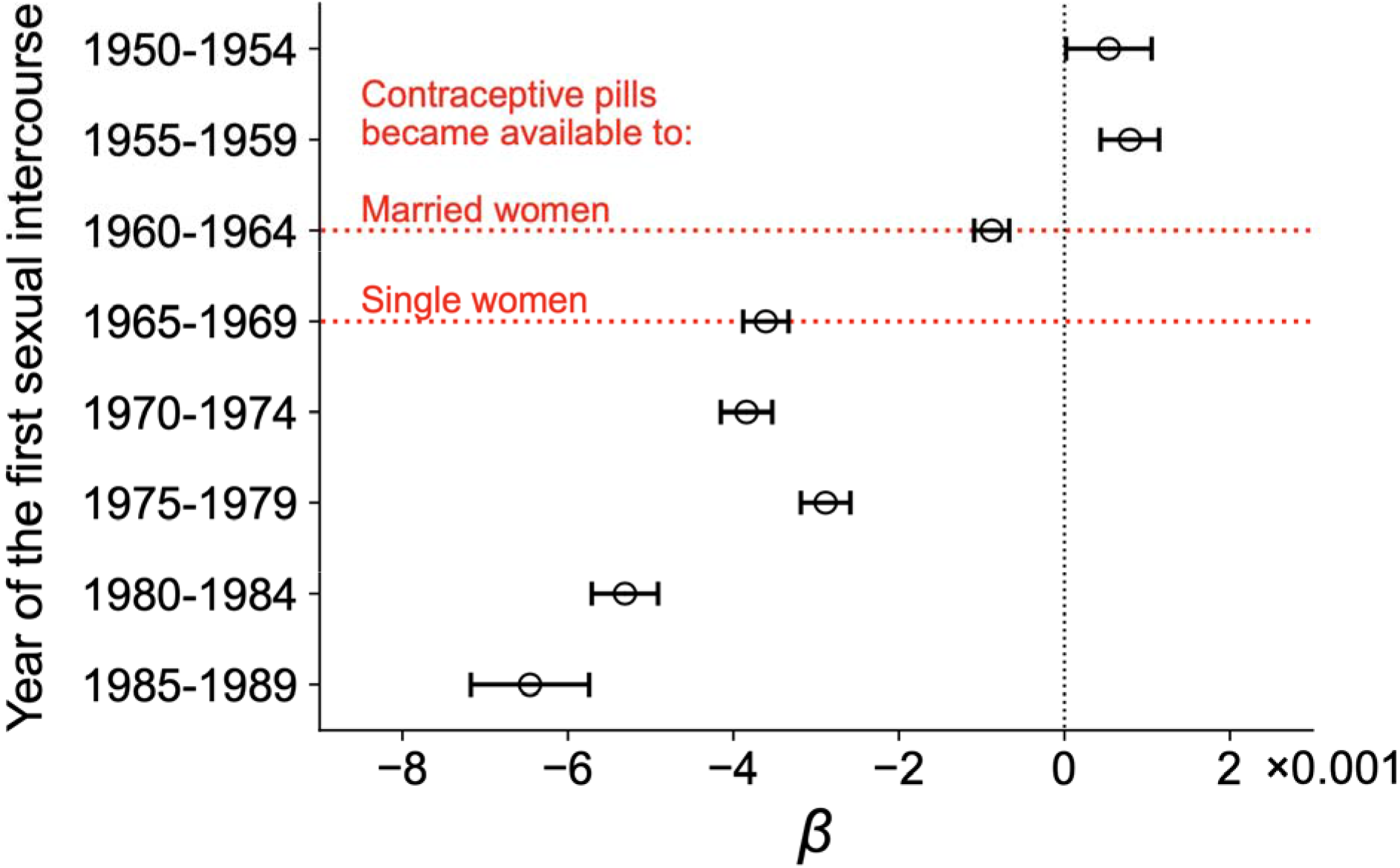
Phenotypic effect (*β*) of the number of sexual partners on the number of children in OSB individuals in each of eight 5-year cohorts. Error bar represents the standard error.

The observation in Fig. 1 prompted the question of whether the genetic correlation between SSB and the number of children among all individuals has become negative since the availability of oral contraceptives in the 1960s. Because an increase in contraception could alter the genetic architecture of fecundity (but not that of SSB), we acquired GWAS summary statistics of the number of children for each of four UKB cohorts classified by the decade in which a participant’s first sexual intercourse occurred. Here, four instead of eight cohorts were considered because estimating the genetic correlation requires SSB individuals, whose numbers are relatively low. Interestingly, the genetic correlation was significantly positive for the 1950s cohort (*r_g_* = 0.73, *P* = 0.003; **Fig. 2**), indicating that SSB-associated alleles overall conferred a positive reproductive effect in the 1950s. Note that this does not necessarily mean that SSB-associated alleles were positively selected, because these alleles could have detrimental non-reproductive effects such as increasing mortality (14) that offset their reproductive benefits. The genetic correlation became slightly but not significantly negative in the cohorts of the 1960s and 1970s but became significantly negative in the cohort of the 1980s (*r*_g_ = −0.33, *P* = 4×10^-4^; **Fig. 2**). That is, SSB-associated alleles are overall reproductively detrimental in the contemporary British population.

**Fig. 2.**
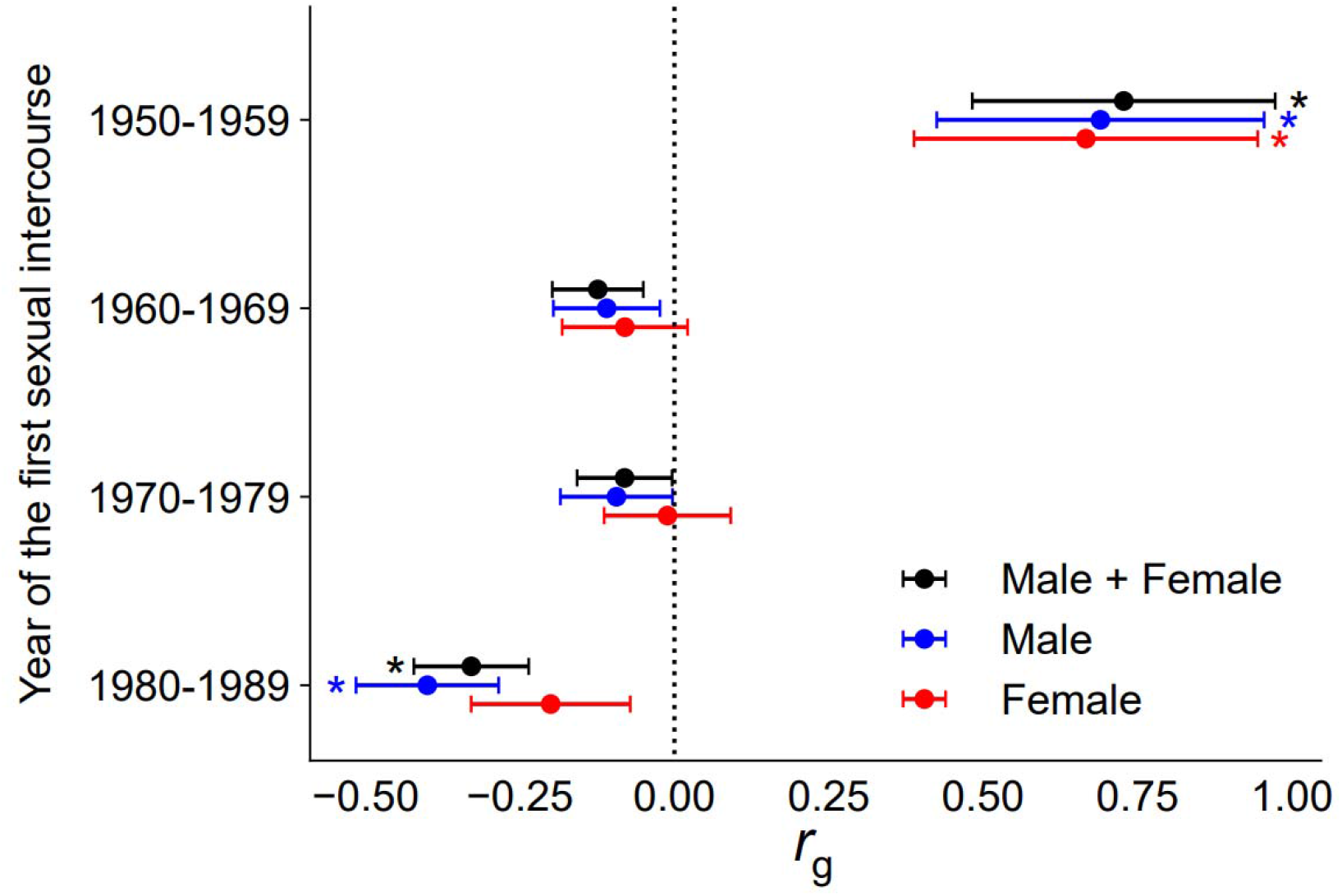
Genetic correlation (*r*_g_) between SSB and the number of children in each of four decades for male (blue), female (red), or both groups of participants (black). Error bars represent standa**rd** errors, whereas asterisks indicate *P* < 0.05.

In summary, if the primary mechanism responsible for the past evolutionary maintenance of human SSB-associated alleles was their offering of reproductive benefits to OSB individuals via boosting their number of sexual partners, this mechanism is no longer at work in modern societies and probably has been gone for at least a half century due to the widespread use of contraception. Remarkably, the number of sexual partners now negatively predicts the number of children in OSB individuals, probably because, given the fixed amount of time that a person has, there is a tradeoff between the time for sexual relationships and childcare once they can be separated by contraception. Hence, the overall impact of SSB-associated alleles on reproduction should now be negative. Indeed, SSB is now negatively genetically correlated with the number of children among all individuals (**Fig. 2**). Therefore, we predict that SSB-associated alleles will gradually decline in their frequencies unless new maintenance mechanisms appear.

Notwithstanding, our prediction does not necessarily imply that SSB will decline, because SSB is also influenced by environmental factors. In fact, the proportion of European ancestry participants in the UKB reporting SSB has generally risen over the four cohorts described in Fig. 2 (2.8%, 2.2%, 3.9%, and 5.4%, respectively), likely attributable to a growing openness of the society toward SSB that could increase the probability of performing and/or the probability of reporting SSB (15).

## Materials and Methods

We analyzed European ancestry individuals in the UK Biobank. GWAS, genetic correlation analysis, and phenotypic regression are conducted. See supporting information (SI Appendix) for additional details.

## Data, Materials, and Software Availability

The genetic and phenotypic data are provided by the UK Biobank. LDSC was used to calculate genetic correlation and can be downloaded at https://github.com/bulik/ldsc. REGENIE was used to perform GWAS and can be downloaded at https://rgcgithub.github.io/regenie/. Other data are included in the article and/or SI Appendix.

## Acknowledgments

We thank J. Li for discussion and E. Long, D. Jiang, and P. Chen for valuable comments. This work was supported by the U.S. National Institutes of Health (grant R35GM139484 to J.Z.).

## Author contributions

S.S. and J.Z. designed the study and wrote the paper; S.S. performed the analysis.

## Competing interests

The authors declare no competing interests.

## Supporting Information

Appendix 1 (PDF)

Dataset S1 (XLSX)

Dataset S2 (XLSX)

Dataset S3 (XLSX)

Dataset S4 (XLSX)

Dataset S5 (XLSX)

## Notes

### Competing Interest Statement

The authors have declared no competing interest.

