## Appendix 1 for "Contraception ends the genetic maintenance of human same-sex sexual behavior"

**This PDF file includes:**

Supporting text: Extended Methods

Legends for Datasets S1 to S6

SI References

**Other supporting materials for this manuscript include the following:**

Datasets S1 to S6

Supporting Information Text

Extended methods

**UKB sample screening.** The current study was approved by the UKB (reference no. 48678). We generally followed the sample screening process previously described (1). To avoid the influence of population stratification, we focused our analysis on the European ancestry population in the UKB (2). We applied the K-means clustering algorithm on the first four genetic principal components precalculated by the UKB (data-field 22009). The resulting four clusters (K = 4) were visually inspected to identify the cluster corresponding to the European ancestry population that contains all individuals labeled as “white British” by the UKB. We further excluded individuals who do not self-report as “white” (data-field 21000) and individuals whose genotype missing rate is greater than 0.02. The resultant 452,557 individuals were both genetically determined and self-identified as “white”, from which 374,585 participants answered questions on sexual partners and children; these answers allowed the identification of 358,861 OSB individuals. GWAS was conducted with appropriate sample set according to the trait of interest.

**UKB variant screening.** We generally followed the variant screening process previously described (1). We excluded single nucleotide polymorphisms (SNPs) that satisfy any of the following conditions: (1) minor allele frequency (MAF) < 1%, (2) imputation quality is low (INFO < 0.8), (3) genotyping missing rate > 1%, and (4) Hardy-Weinberg equilibrium (HWE) is rejected at P < 10−10. In the end, 9,371,426 SNPs were used in the analysis, including SNPs on the X chromosome.

**UKB phenotypes investigated.** The phenotype of same-sex sexual behavior (SSB) in the UKB was acquired from the answer to the question “Have you ever had sexual intercourse with someone of the same sex?” (data-field 2159). The information on the number of children was obtained from the answers to the question “How many children have you fathered?” (data-field 2405) and “How many children have you given birth to? (Please include live births only)” (data-field 2734). The information on the number of sexual partners of an OSB individual was obtained from the answer to the question “About how many sexual partners have you had in your lifetime?” (data-field 2149). Individuals who refused to answer a question were excluded from the analysis on the relevant phenotype.

**Phenotypic regression.** Among the 452,557 European ancestry samples, 374,585 provided valid information about the number of sexual partners (data-field 2149) and the number of children (data-field 2405 or data-field 2734), and we focused on 358,861 OSB individuals. Linear regression of the number of children on the number of sexual partners in OSB individuals was performed with genetic sex (data-field 22001), age (data-field 21022), age squared, and the first ten genetic principal components (data-field 22009) used as covariates. Socioeconomic confounding factors were also controlled by including household income (data-field 738), deprivation index (data-field 26410), number of years of education (derived from data-field 6138 according to ref.), year of first sexual intercourse (derived from data-field 34 and data-field 2139), and their squared terms as covariates.

**GWAS.** To perform GWAS, we applied the state-of-the-art software REGENIE (3). REGENIE analysis consists of two steps. The first step fits a whole genome regression model for trait predictions based on high-quality genotyped SNPs using the leave one chromosome out (LOCO) scheme.  In the second step, the phenotypic predictions attained from the first step are used as offsets and GWAS is conducted for all SNPs—both genotyped and imputed—using standard linear regressions.  For the first step, we used all 595,864 genotyped SNPs satisfying the following: (1) minor allele frequency (MAF) > 1%, (2) minor allele count (MAC) > 100, (3) genotyping missing rate < 1%, and (4) Hardy-Weinberg equilibrium (HWE) test *P* > 10^−10^. For the second step, we used the previously described screened European ancestry individuals and variants (9,371,426). The same covariates were used in regression models for both steps, including age, age squared, genetic sex, and the first ten genetic principal components.

**Genetic correlation.** We used cross-trait linkage disequilibrium (LD) score regression based on GWAS summary statistics to estimate the genetic correlation between traits (4). Briefly, the genetic correlation is estimated by calculating the slope of the regression where the products of z-scores resulting from GWAS of two traits are regressed upon the LD score. We used the software LDSC (5) and followed its instructions (https://github.com/bulik/ldsc) to calculate genetic correlations.

**UKB cohorts by year of first sexual intercourse.** We divided OSB individuals of European ancestry into eight 5-year cohorts according to their year of first sexual intercourse (derived from UKB data-field 34 and data-field 2139): 1950-1954, 1955-1959, and so on. We performed phenotypic regression of the number of children on the number of sexual partners in OSB individuals in each cohort with the same method described in the “phenotypic regression” section. The effect size (*β*) of the number of sexual partners and its statistical significance were reported (**Fig. 1**). For cohort-stratified genetic correlation between SSB and number of children, we used four cohorts (1950-1959, 1960-1969, 1970-1979, and 1980-1989) due to the relatively small numbers of SSB individuals in the data. We performed the genetic correlation as described in the “genetic correlation” section (**Fig. 2**).

**Validation using multiple recruitment centers.** The UKB has 22 recruitment centers that recruited participants from different geographic areas in the UK from 2006 to 2010. Therefore, cohorts recruited by different centers with the same protocol can be viewed as independent replicates. We replicated the phenotypic regression, GWAS, and genetic correlation analyses for each of the 19 recruitment centers with at least 10,000 participants.

**Potential influence of cross-trait assortative mating on genetic correlation.** It was reported that cross-trait assortative mating could generate genetic correlations between traits that are not due to their shared (or linked) genetic variants (6). For example, assortative mating between individuals with relatively high numbers of sexual partners and those with relatively low numbers of children could generate a negative genetic correlation between the number of sexual partners and the number of children even when the two traits share neither genetic variants nor linked genetic variants. Nevertheless, even in the above scenario, a high number of sexual partners genetically predicts a low number of children because alleles associated with more sexual partners and alleles associated with fewer children tend to be in the same individual due to assortative mating. Hence, potential cross-trait assortative mating does not compromise our interpretation of genetic correlation in the present study.

**Potential reasons why Zietsch *et al.* observed a positive genetic correlation between SSB and number of children.** We detected no significant genetic correlation between SSB and the number of children in OSB individuals (**Dataset S5**). However, Zietsch *et al.* (7) found that SSB is genetically significantly positively correlated with number of children in males (their Table S2). There are several reasons why their results differed from ours. First, Zietsch *et al.* used previously published GWAS summary statistics for SSB (1) and number of children (8) that were based on European ancestry individuals from the UK (mainly the UKB), other European countries (*e.g.*, deCODE), and the US (*e.g.*, 23andMe), while our analysis exclusively used European ancestry individuals from the UKB. The difference is that our sample contains a higher fraction of British ancestry individuals than Zietsch *et al.*’s sample. Second, the social environment can differ between countries, which might influence sexual and reproductive behaviors. For example, US teenagers have higher pregnancy and reproduction rates relative to those of British teenagers, and US women tend not to use contraception compared with British women (9). These differences can influence the genetic correlation between SSB and number of children. Third, Zietsch *et al.* calculated the genetic correlation between SSB and number of children in the whole population including both SSB and OSB individuals, while our Dataset S5 presented the genetic correlation between SSB and number of children in OSB individuals. Our Fig. 2 made it clear that the genetic correlation between SSB and number of children in the whole population was significantly positive in the 1950s cohort but became negative in the 1960s and 1970s cohorts and significantly negative in the 1980s cohort.

Dataset S1 (separate file). Phenotypic effect (*β*) of the number of sexual partners on the number of children in OSB individuals recruited by different UKB recruitment centers.

Dataset S2 (separate file). Genetic correlation (*r_g_*) between the number of sexual partners (NSP) and the number of children (NC) in OSB individuals recruited by different UKB recruitment centers.

Dataset S3 (separate file). Distribution of the year of the first sexual intercourse among UKB participants of European ancestry.

Dataset S4 (separate file). Genetic correlation (*r_g_*) between SSB and the number of sexual partners among OSB individuals.

Dataset S5 (separate file). Genetic correlation (*r_g_*) between SSB and the number of children among OSB individuals.
